# Heritability in friendship networks

**DOI:** 10.1101/2019.12.31.891598

**Authors:** Michael Neugart, Selen Yildirim

**Affiliations:** Department of Law and Economics, Technische Universität Darmstadt, Darmstadt, Germany; Graduate School of Economics, Finance and Management, Goethe-Universität Frankfurt, Germany

**Keywords:** social networks, twins, behavioral genetics

## Abstract

Friendship networks account for a large part of an individual’s economic success or failure in life. Using data from the German TwinLife study, we explore, within a classical twin design, to which extent friendship networks are related to genes. We find a substantial heritability component in twins’ network sizes and network homophily, but not in twins’ network closeness. Addressing indirect ways in which genes could influence network characteristics, we do not find evidence that shared hobbies affects networks.

## 1 Introduction

Friendship networks exhibit sizable variation. Some people have numerous friends, others rely only on a few. Some people’s friends know each other, other friendship networks hardly overlap. Moreover, some people have heterogeneous friends, while in other friendship networks similar people flock together. To understand this variation in friendship networks, which very often determines an individual’s economic success or failure in life, scholars have developed various network formation models [1], often disregarding the effects of an individual’s traits.

The genetic influence is nowadays widely accepted for many traits [2]. There is, however, little work on the association of genes with an individual’s overall network characteristics [3]. We explore to which extent friendship networks are related to genotypes. Using a classical twin design, we detect the share of observed variation in network size, closeness, and homophily associated with genes.

There is also limited knowledge on the channels through which genes may affect network characteristics. It is conceivable that an individual’s network is not directly associated with an individual’s genetic disposition. One may also surmise that network traits are the result of genetically determined shared activities of twins [4]. We analyze twin data which allows us to estimate the variance components of the network traits associated with genes by twins who share hobbies and who do not share hobbies. Thus we are able to identify whether the heritability of friendship networks is intermediated by a selection of twins into same hobbies and interests.

## 2 Materials and Methods

### 2.1 Participants

Our data come from the TwinLife project, a multi-disciplinary twin study conducted in Germany [5]. The survey began in 2014 and comprises four cohorts (1: 2009/2010, 2: 2003/2040, 3: 1997/1998, 4: 1990-1993). Zygosity is determined on questionnaires. Adult twins (or the parents of twins under the age of 16) report physiological characteristics, eye and skin color, frequency of confusion of the twins by others etc. In addition, there are questions on self-rated zygosity, and on twin specific issues like hobbies or shared interests. Our analysis draws on information of 4,136 same-sex twins. 1,990 are identical twins (monozygotic, MZ) and 2,146 are fraternal twins (dizygotic, DZ) twins. Among the MZ twins 772 are male, and among the DZ twins 918 are male.

To explore the representativeness of the TwinLife data, Lang and Kottwitz [6] compare the distributions of the key socio-demographic characteristics of the TwinLife sample with those found in the German Microcensus sample by creating a proxy-twin and multiple-child household. Their analyses show that most socio-demographic indicators are similar for proxy-twin and multiple-child households between the two data sets.

### 2.2 Measures

We are interested in the genetic determinants of twins’ friendship networks. The data allows us to construct four different measures of friendship network characteristics: network size, network closeness, and network homophily by age and by gender. Table 1 shows the questions to the twins on which the coding of the variables is based.

**Table 1.** Definition of variables.

| Variable | Description |
| --- | --- |
| Network size | Network size measures the number of close friends. This does not include family members or Facebook friends. |
| Network closeness | The variable is created using the answer to the survey question “When my friends celebrate their birthdays there are usually many people that I don’t know.”. Answers to this question range from one (totally disagree) to four (totally agree). Network closeness is a binary indicator which is assigned the value one if twins answer the survey question with a one or a two, and is assigned the value zero otherwise. |
| Homophily:<br>Similar age | Homophily: Similar age is a binary indicator that is assigned the value one if the age difference (in years) between the twin and all three important persons is less than or equal to one, and is assigned the value zero otherwise. |
| Homophily:<br>Same-sex | Homophily: Same-sex is a binary indicator which is assigned the value one if all three important persons and the twin are of same sex, and is assigned the value zero otherwise. |
| Under 18 | Under 18 is a binary variable which is assigned the value one if twins are younger than 18 years, and is assigned the value zero otherwise. |
| East Germany | The federal state indicator equals one, if the residence is in a federal state in East Germany and is assigned the value zero otherwise. |
| Shared<br>interests/hobbies | The survey includes a question on whether the twins (currently) share the same hobbies and interests. The possible answers to this question are “1: rather yes”, “2: rather no”, and “3: in the past yes, now no longer.”. Shared interests/hobbies is a binary indicator which is assigned the value one if the answer to the question is one or three, and is assigned the value zero otherwise. |

The median number of friends twins have (network size) is seven and equal for MZ and DZ twins, see Table 2. About 80% of the twins typically know the friends of their friends (network closeness). In relation to network homophily: 52% of the MZ twins and 58% of the DZ twins have three most important friends with an age difference of at most one year, respectively, and 20% of the MZ twins and 18% of the DZ twins have three most important friends of the same gender. The TwinLife project focuses on adolescents which is reflected in the median age of 17 years, and a share of 57% of twins in the sample which is under 18 years old. 16% of the twins reside in East Germany. There is information on whether twins share hobbies or interests for 2,946 twins. Among the MZ twins, 97% state to have or to have had shared hobbies. Among the DZ twins 79% state to have or to have had shared hobbies.

**Table 2.**
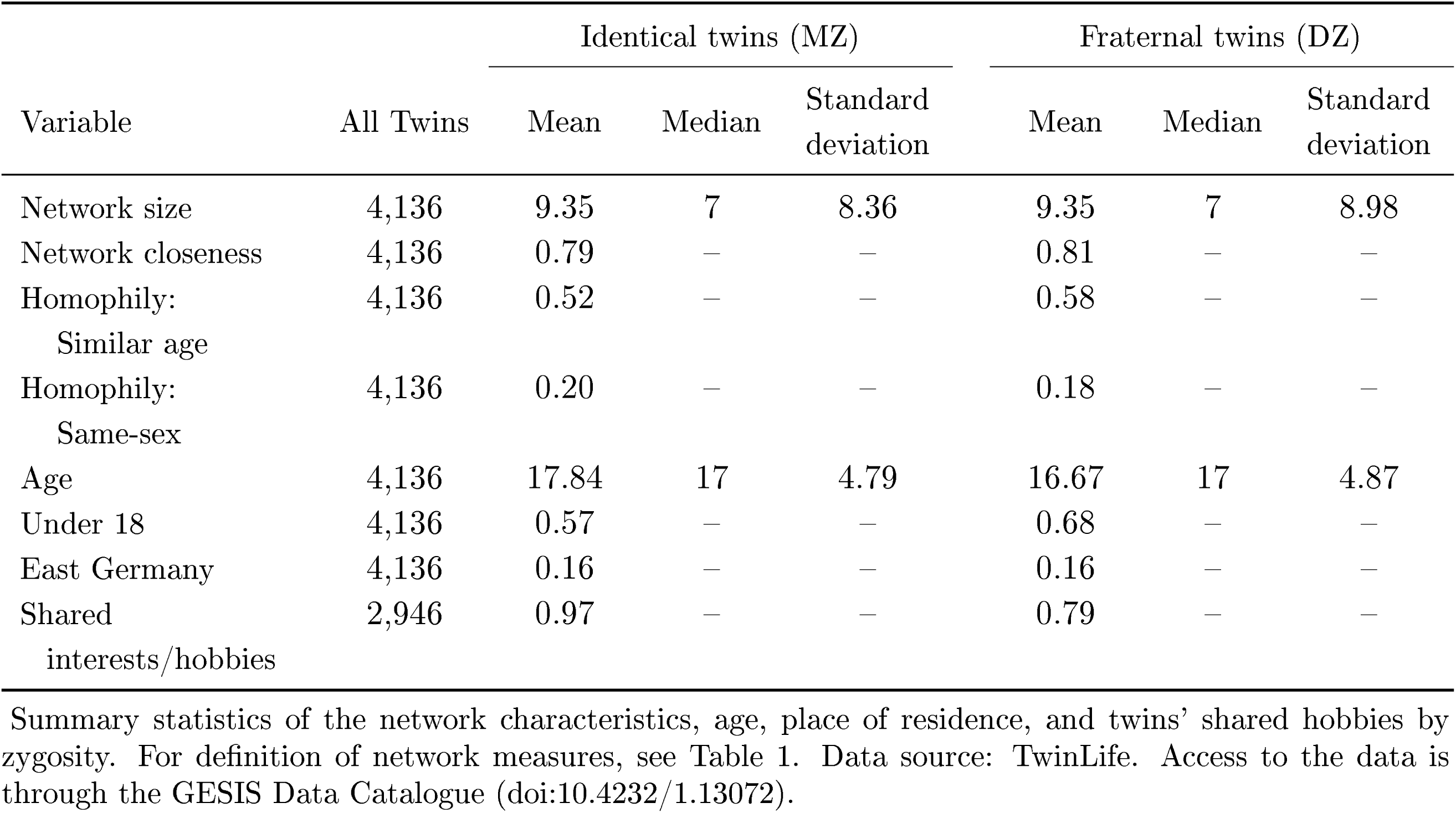
Network characteristics and control variables by zygosity.

### 2.3 Analysis

To test the hypothesis that genes play a role in friendship networks, we use the classical twin design [7]. The heritability of a trait is measured by comparing the trait similarity of MZ twins, who share 100% of their genes, and the trait similarity of DZ twins, who share only 50% of their genes – on average. Under the assumptions of the classical twin design, MZ twins should be more similar than DZ twins if genetic variation is a factor contributing to a given trait. The design is accepted as one of the most powerful study designs for decomposing the relative contribution of genes and the environment to human behavior and traits.

More specifically, biometrical genetic theory enables us to write structural equations relating the observed behavior of twins to their underlying genotypes and environments [8]. The observed behavior, *y*_*ij*_, of identical and fraternal twins can be modeled for twin *i* (1 or 2) of pair *j* as a function of three unobservable effects, namely an additive genetic (A), a common environment (C), and a unique environment (E) component. The model, which is typically referred to as the “ACE”-model, can be written as:

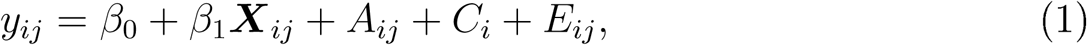

where *β*_*o*_ is an intercept term and *β*_1_ captures the effects of the covariates *X*_*ij*_. *A*_*ij*_ ∼ *N* (0, *σ*_*A*_) is a genetic component, *C*_*i*_ ∼ *N* (0, *σ*_*C*_) a common environment component, and *E*_*ij*_ ∼ *N* (0, *σ*_*E*_) is a unique environment component.

## 3 Results

### 3.1 ACE decomposition

Fig 1 shows the results of an ACE-decomposition on the four network traits for the full sample (black bars) and a restricted sample (blue bars). For the full sample, 54% of the variation in observed network size can be attributed to genes. Network closeness is not related to heritability. Homophily, defined as a network of friends of the same gender, and homophily, defined as a network of friends of similar age, display almost the same share of heritability, i.e. 42% and 43%, respectively. The common environment components are not statistically different from zero for network size, network closeness, and network homophily based on age, but is positive for network homophily based on gender. The rest of the observed variation in all four traits falls on an idiosyncratic environment.

**Figure 1.**
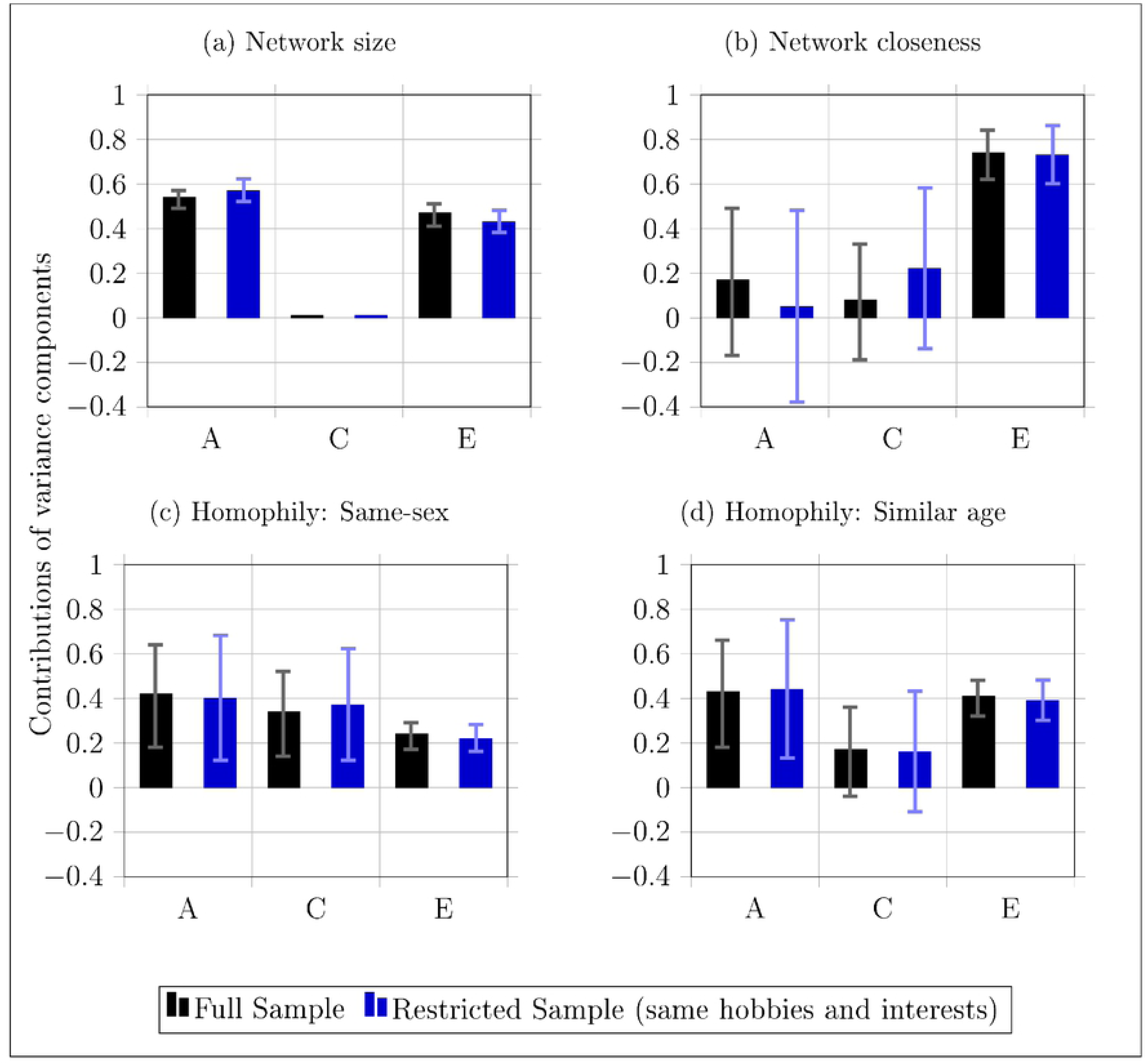
ACE decomposition. The above panels report the results from maximum likelihood estimations of behavioral genetic variance decomposition models. The full sample, represented by black bars, consists or 4,136 twins. The restricted sample, represented by blue bars, consists of 2,608 twins who share hobbies and interests. The variance of network characteristics of twins are partitioned into additive genetic (A), common environment (C), and unique environment (E) components, while controlling for age of twins, sex, and place of residence. Standard errors are clustered at family level. The coefficients and 95% confidence intervals are displayed for each model. We report zero when the non-negativity constraint is binding for a variance component.

### 3.2 Mechanism

Our estimates of the ACE components of the four network traits indicate that genes play a key role in people’s friendship networks. Jackson [4] conjectures that the genes, however, may not only be directly related to the observed variation in people’s friendship networks. He hypothesizes that other potentially heritable traits might indirectly influence network characteristics. In particular, heritable traits such as physical or mental capabilities may affect twins’ leisure activities. This could then lead to MZ twins having a greater overlap in their activities than DZ twins, and, therefore, a greater overlap in their common environment. For example, twins may have a preference for and the skills needed to excel at a team sport like soccer. In contrast to twins who are leaning towards individual sports, twins conducting team sports would very likely have larger, more homophilous, and closer networks.

Using information on twins’ shared hobbies and interests, we investigate whether such an indirect channel exists. To this end, we restrict the sample to twins who report having shared hobbies and interests, and re-run our ACE-decomposition. If an indirect channel exists, we expect a lower A-component in the restricted sample because a possibly genetically driven selection into shared activities is addressed by the stratification of the sample.

The blue bars in Fig 1 display the ACE-components estimated for twins who share hobbies and interests. Restricting the analysis to MZ and DZ twins with shared hobbies and interests reduces the sample to 2,608 observations. As can be seen from Fig 1, the ACE-components do not differ from the components estimated on the full sample. Thus we conclude that genetically determined selection into shared activities does not indirectly influence the network characteristics in our sample of twins.

### 3.3 Equal treatment of parents

A common critique of twin studies is that, in contrast to the underlying assumption in the classical twin design, they usually cannot take into account unequal treatment of identical twins and fraternal twins by the parents [9]. We attempt to remedy concerns in relation to the equal environment assumption by making use of a large set of questions posed to the twins in the TwinLife study about how their parents treated them.

In total, we have 13 questions that relate to parenting style. For example, one such question to the twin is on how many times the father or mother “… shows you that s/he likes you.” We use this information to compare the parenting styles of fathers and mothers who either have MZ or DZ twins. A detailed description of all questions related to parenting styles is in the supplementary material S1 Table.

In only three out of the 13 answers to the questions on the parenting style do we find significant differences in how the father *and* mother have been treating MZ and DZ twins. Further we conduct an analysis in which we restrict the sample to the twin pairs who responded to the parenting style question with the same answers, or the difference between answers is less than two on a scale between zero and four. For these twins, who report being treated equally by their parents, we estimate ACE-components as shown in Fig 2. Apparently, these estimates are close to those found in the baseline model. They mostly lie within the confidence intervals estimated for the full sample. We interpret this as evidence in support of an un-biased estimate of the heritability component when we do the variance decomposition on the full sample.

**Figure 2.**
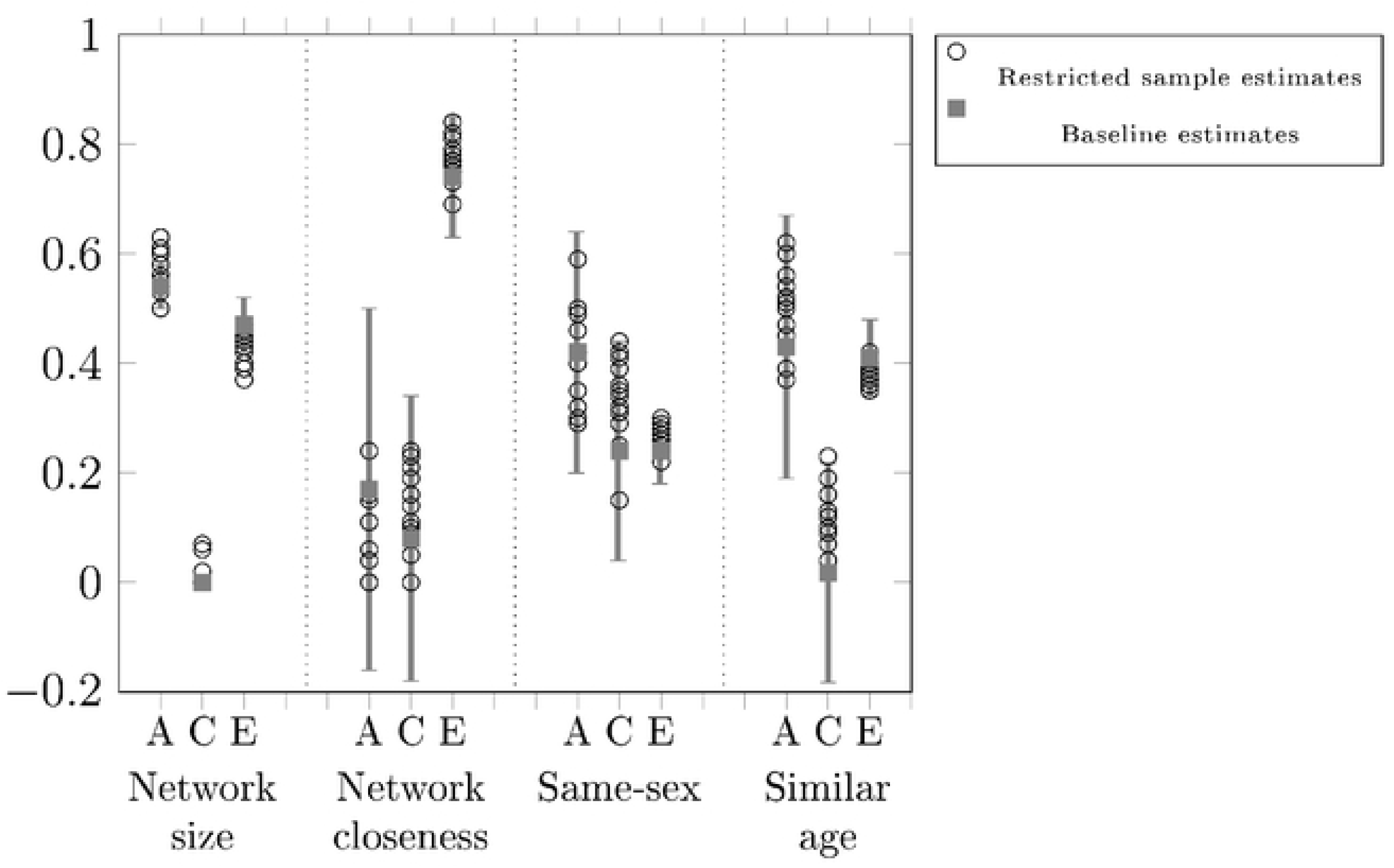
Robustness – ACE decomposition for twins treated equally by their parents. reports the results from maximum likelihood estimations of behavioral genetic variance decomposition models. The variance of network characteristics of twins are partitioned into additive genetic (A), common environment (C), and unique environment (E) components, while controlling for age of twins, sex, and place of residence. Standard errors are clustered at family level. The estimates from the baseline model, c.f. Figure 1 (full sample), with 95% confidence intervals are displayed for each netwrok characteristic. The baseline estimates are compared to the results of “ACE”-models using samples of twins equally treated by their parents for each of the 13 parenting style questions.

## 4 Discussion

Using a classical twin design, we explore heritability in friendship network characteristics. We find a considerable genetic component in twins’ network size (54%) and network homophily (42% when we measure homophily with same gender and 43% when we measure homophily with similar age), but not in twins’ network closeness. The results are in line with estimates of sizable heritability components in overall network characteristics of an earlier study on 1,100 twins in 142 separate high school friendship networks [10]. In this study, which was based on interviews for the United States National Longitudinal Study of Adolescent Health, the following results were found: genetic components of 46% for an in-degree measure of the network, 47% for a transitivity measure of the network, 29% for a betweenness centrality measure of the network, and a not significant genetic component for an out-degree measure of the network. Not only do our results on the heritability of network characteristics comply with that previous work, our results are also robust with respect to the equal parenting assumption which underlies the classical twin design.

Three of the four network characteristics have a positive heritability component indicating that network traits are associated with genes. It is an open question why variation in network closeness has a zero heritability component. A possible reason could be that network closeness is not related to genetically determined personality traits which facilitate introducing friends to other friends.

What is also novel about our study is that we analyze an indirect channel through which genes may affect network characteristics. We conjecture that MZ twins more likely select themselves into the same hobby than DZ twins, which may affect their friendship networks. We do not find evidence that shared hobbies and interests affects networks. Our measure of shared hobbies and interests is, however, insofar limited as we do not have information on the type of hobbies and interests shared. Hobbies could involve many other people as in team sports or it could be an individual sport with less people being involved. It could be insightful to analyze twin data by type of interest and hobby to elicit whether network measures change. We suspect that, in particular, network closeness would be associated with genes differently as we were able to split samples by type of hobbies, i.e. a team sport and an individual sport.

Our results suggest that in three of the four network measures there is a genetic component. The remaining variation in the network traits falls on a common environment and on an idiosyncratic environment. For three of the four traits the idiosyncratic environment appears to play a more important role than the common environment. Large idiosyncratic components, relative to the common environment, have been reported elsewhere for traits in the social domain [2]. Our study adds to these results that the more salient environmental influences are those idiosyncratic to a child.

Overall, our results motivate to shift greater attention to the development of network formation models that take into account genetic variation in order to better understand how social structures influence an individual’s economic choices.

## Supporting Information

S1 Table. Questions on parenting style.

